# Modelling nonlinear dendritic processing of facilitation in a dragonfly target-tracking neuron

**DOI:** 10.1101/2021.03.24.436732

**Authors:** Bo M. B. Bekkouche, Patrick A. Shoemaker, Joseph M. Fabian, Elisa Rigosi, Steven D. Wiederman, David C. O’Carroll

## Abstract

Dragonflies are highly skilled and successful aerial predators that are even capable of selectively attending to one target within a swarm. Detection and tracking prey is likely to be driven by small target motion detector (STMD) neurons identified from several insect groups. Prior work has shown that dragonfly STMD responses are facilitated by targets moving on a continuous path, enhancing the response gain at the present and predicted future location of targets. In this study, we combined detailed morphological data with computational modelling to test whether a combination of dendritic morphology combined with the nonlinear properties of NMDA receptors could explain these observations. We developed a hybrid neuronal model of neurons within the dragonfly optic lobe, which integrates numerical and morphological components. The model was able to generate potent facilitation for targets moving on continuous trajectories, including a localized spotlight of maximal sensitivity close to the last seen target location, as also measured during *in vivo* recordings. The model did not, however, include a mechanism capable of producing a traveling or spreading wave of facilitation. Our data support a strong role for the high dendritic density seen in the dragonfly neuron in enhancing non-linear facilitation. An alternative model based on morphology of an unrelated type of motion processing neuron from a dipteran fly required more than 3 times higher synaptic gain in order to elicit similar levels of facilitation, despite having only 20% fewer synapses. Our data supports a potential role for NMDA receptors in target tracking and also demonstrates the feasibility of combining biologically plausible dendritic computations with more abstract computational models for basic processing as used in earlier studies.

## 1 Introduction

Imagine a dragonfly hungry for breakfast, flying out over a small lake in the morning and searching for small prey to catch. As the dragonfly approaches a swarm of flies, it focuses on one whilst ignoring the others, before pursuing and catching its prey mid-air. Such pursuits typically succeed over 95% of the time (Olberg et al., 2000). The selective attention required to mediate this behavior (Wiederman and O’Carroll, 2013) is aligned with two fundamentally distinct computational principles. The first is a winner-take-all network, required to ignore distractors (Shoemaker, 2015).

The second is neuronal facilitation, which boosts the gain of the response for the attended target as it moves along continuous trajectories, anticipating its future path and boosting its intrinsic salience when seen against complex backgrounds of visual clutter (Nordström et al., 2011; Dunbier et al., 2012; Bagheri et al., 2017; Wiederman et al., 2017).

In the insect brain, these computations are believed to take place in the medulla and lobula of the optic lobes, either presynaptic to, or within the dendrites of small target motion detectors (STMDs), a neuron type described from the insect lobula and midbrain (e.g. **Figure 1**). STMDs respond selectively to small moving targets, ignoring larger features (O’Carroll, 1993; Nordström and O’Carroll, 2009). STMD neurons are a diverse group, including subtypes that are both afferent or efferent, and with a wide range of receptive field size, location and direction selectivity. For example, receptive fields may be as small as just a few degrees of visual angle within a subtype termed small field STMDs (SF-STMDs) which are found in both dipteran flies and dragonflies (Barnett et al., 2007; Wiederman et al., 2017). Other neurons such as the identified binocular dragonfly STMD neuron BSTMD1, may give excitatory responses to stimuli presented anywhere in the dorso-frontal visual fields of either eye (Dunbier et al., 2012). Complex excitatory and inhibitory interactions between both visual fields are mediated by heterolateral and centrifugal STMD neurons such as the identified cell CSTMD1. This neuron type has been identified in both dipteran flies and dragonflies (Nordström et al., 2006; Geurten et al., 2007), and has its inputs in the midbrain, with extensive arborizations in the contralateral lobula.

**Figure 1.**
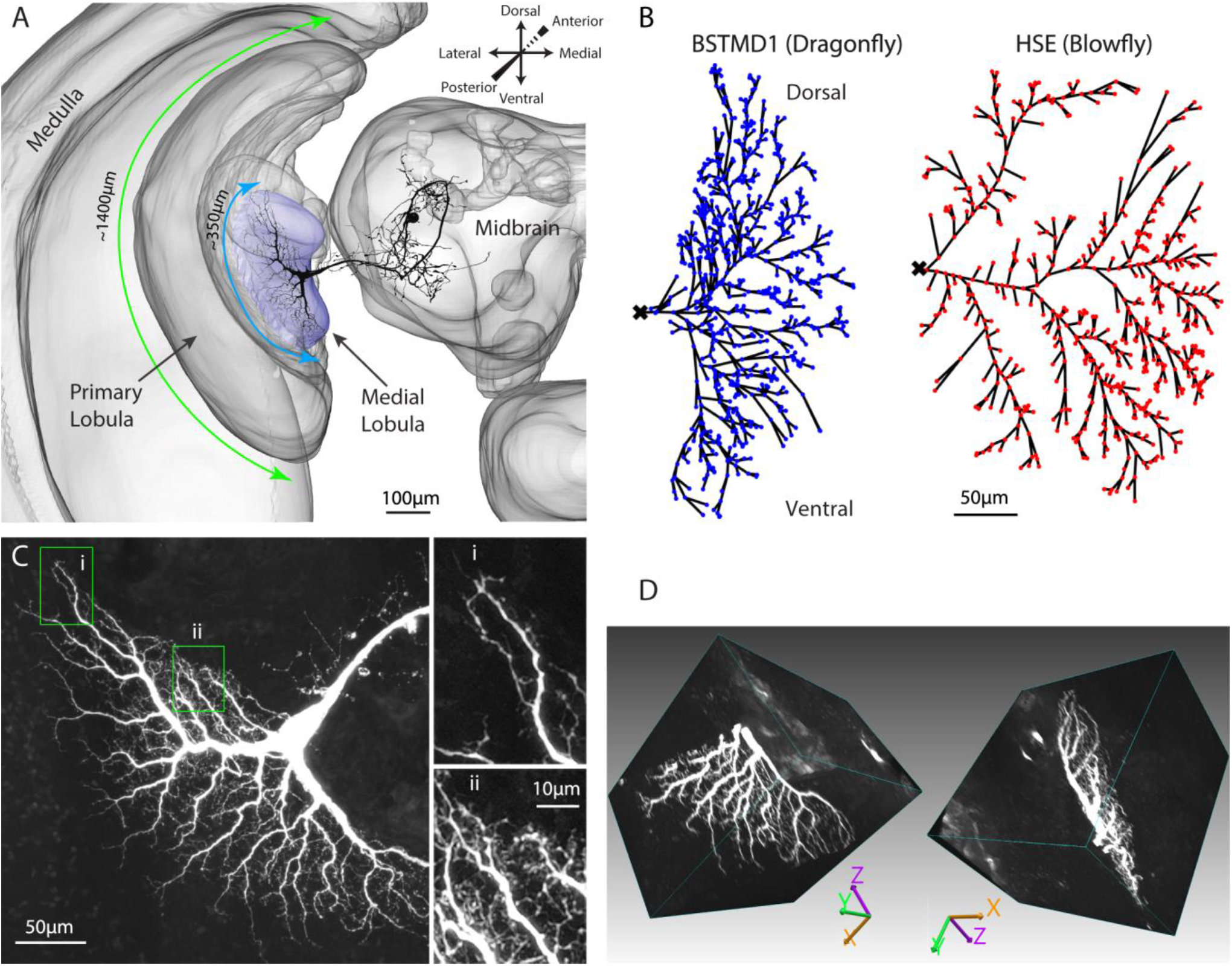
Anatomy of the dragonfly brain and dendritic arborisations of BSTMD1. **(A)**: A 3D reconstruction of the brain showing the location and scale of the BSTMD1 neuron, with its main input arborization in the medial lobula. Green and blue arrows denote the approximate span of the outer part of the primary lobula and the medial lobula, respectively. **(B)** Z-projections from reconstructed 3D anatomical models for the main input arborization of the BSTMD1 neuron (left) and of an HSE neuron from the blowfly (right). The BSTMD1 model was based on data from the present study, while the HSE model used data downloaded from neuromorpho.org (Ascoli et al., 2007; Cuntz et al., 2008). The dots indicate the location at which an NMDA receptors (or double exponential receptors) were placed within the arborizations in the multicompartment model. The ‘x’ marks the junction of the dendritic trees with the main arborization on the medial side of the tree, and also the location from which the membrane potential was estimated in subsequent multicompartment modeling. **(C)** Z-projection (maximum intensity) of a confocal image volume showing the main BSTMD1 arborisation. Insets show higher magnification views of dense fine dendrites, featuring both dendritic spines and ‘blebs’ that lie beyond the resolution of the anatomical model in B. **(D)** Views of a rendered 3D volume image (opaque maximum intensity projection) from the ventral half of the arborization and rotated by different amounts. The left view approximately matches the Z-projection in C.

Because of its expression of stimulus history-dependent predictive gain enhancement, and the impressive ability to select and attend to single targets amidst alternative distracters, CSTMD1 and its presumptive inputs from neurons such as BSTMD1 have become important models for studying mechanisms of target selection and attention (Wiederman and O’Carroll, 2013; Wiederman et al., 2017; Lancer et al., 2019). The underlying neural mechanism of facilitation as observed in dragonfly STMDs is not known. However, a non-linearly amplifying receptor subtype, such as the N-methyl-D-aspartate receptor (NMDA) receptor expressed in glutamatergic synapses, has been proposed as a candidate mechanism (Shoemaker, 2011; Bekkouche et al., 2017). When an NMDA receptor binds glutamate, the ion channel opens for sodium and calcium ions to pass into the neuron, but this occurs only if the neuron is already partially depolarized (required to displace magnesium ions that block the channel at hyperpolarized potentials). In this way, NMDA receptors respond supralinearly to input: the more depolarized the local membrane potential becomes by synaptic currents, the greater the NMDA conductance. This leads to an enhanced response (facilitation), at least up to a certain turning point, after which it behaves more linearly (Xia and Chiang, 2009). We do not yet know how widely the NMDA receptor is expressed in dragonfly brains. However, the *Drosphila melanogaster* NMDA homologue is widely expressed in the insect medulla and lobula (optic lobe) (Xia et al., 2005; Wu et al., 2007; Davis et al., 2020).

While direct evidence has yet to be obtained, we here propose that NMDA synapses may be expressed in the input dendrites of dragonfly STMD neurons. Prior attempts to model this phenomenon as a possible basis for facilitation (Shoemaker, 2011) used the dendritic morphology of a neuron type in which facilitation has not been observed physiologically, a horizontal system (HS) neuron from the lobula plate of a dipteran fly, based on downloadable neuron reconstructions (Ascoli et al., 2007; Cuntz et al., 2008). This was due mainly to a lack of detailed anatomical data at that time for the input dendritic arborizations of suitable STMD neurons. Although this demonstrated that NMDARs can, in principle, produce response time courses for targets moving along long trajectories that resemble those seen in dragonfly STMDs, these models may have missed key network processing properties due to differences between the anatomy between HS neurons and the specific distribution of synaptic zones along the dendritic morphology of dragonfly STMDs. More recently, we developed a modelling framework allowing us to compare the responses of fly HS neurons to those based on newly obtained data for dragonfly STMD neurons (Bekkouche et al., 2017).

In the present study, we further developed this computational approach to investigate the possible role for NMDARs in the facilitation of dragonfly STMD neurons, as introduced in our previous work (Shoemaker, 2011; Bekkouche et al., 2017). Here we focus on the facilitation of responses to stimuli moving along long trajectories in the dragonfly BSTMD1 neuron, believed to play a key role in the dragonfly visual selective attention mechanism (Dunbier et al., 2012). BSTMD1 responds to visual stimuli presented in either visual hemifield, but with different response characteristics. When recording in the lobula from the thick axon near the junction of input dendrites (**Figure 1A**) and the visual stimulus is presented on the ipsilateral side, the neuron gives mixed-mode responses, with spikes riding on a pronounced graded depolarization. Contralateral stimuli elicit larger spikes that ride on a hyperpolarizing graded response, suggestive of complex binocular interactions mediated by the extensive processes of this neuron within the central brain (Dunbier et al., 2012).

From initial anatomical data, Dunbier et al. (2012) proposed that BSTMD1 may make and receive bi-directional excitatory synapses with CSTMD1 and may be involved in attentional modulation of the latter as targets cross the midline between visual hemispheres. In this paper we do not consider these complex binocular interactions. Rather, since both ipsilateral and contralateral stimuli induce strong facilitation in this neuron (Dunbier et al., 2012) we used high-quality confocal images of an intracellularly labeled neuron to develop a detailed anatomical model for BSTMD1’s assumed main ipsilateral inputs, a large dendritic tree located in the lobula. We then developed a multicompartment computational model for synaptic integration of NMDARs based on this anatomical model and its presumed dendritic synaptic nodes. This computational model was then coupled to inputs from a bio-inspired computational model for local target selectivity based on temporal correlation of luminance decrements (OFF stimuli) with subsequent increments (ON stimuli) - a characteristic signature for a small dark target moving against the background (Wiederman et al., 2008, 2013). We then mapped a single stimulus input space (a sequence of images) onto this hybrid model to investigate responses to locally presented stimuli before or after long path ‘primer’ stimuli. Our model was able to generate pronounced facilitation, as also measured during *in vivo* recordings from BSTMD1. Despite varying the synaptic gain, we were unable to recruit such strong facilitation using an alternative model based on the anatomy of a fly wide-field motion neuron, suggesting that this property depends on the unique morphology of BSTMD1.

## Methods

### 1.1 Intracellular labelling and anatomical 3D reconstruction

Wild-caught dragonflies (*Hemicordulia tau*) were immobilized with a 1:1 beeswax and rosin mixture and fixed to an articulated magnetic stand with the head tilted forward to access the posterior surface. A hole was cut above the brain to gain access to the lobula and lateral midbrain, and we then penetrated the perineural sheath and recorded intracellularly using hard aluminosilicate micropipettes (OD=1.00, ID=0.58 mm), pulled on a Sutter Instruments P-97 puller. The electrode tip was filled with 4% Lucifer Yellow solution in 0.1M LiCl and backfilled with 0.1M LiCl. Electrodes were placed in the medial portion of the lobula complex and stepped through the brain from posterior to anterior, using a piezoelectric stepper (Marzhauser-Wetzlar PM-10). Intracellular responses were digitized at 5 kHz with a 16-bit A/D converter (National Instruments) for off-line analysis. The identity of BSTMD1 was confirmed by analysis of its response properties to visual stimuli, and its characteristic binocular receptive field, as described previously (Dunbier et al., 2012). The neuron was then injected with lucifer yellow by passing hyperpolarizing current of −2nA for at least 20 minutes.

Following injection, the brain was carefully dissected under phosphate buffered saline (PBS) and then fixed overnight in 4% paraformaldehyde (in PBS) at 4°C. To intensify the lucifer injection, brains were then rinsed (3×10 minutes) with PBS, before permeabilization in 80/20 DMSO/Methanol solution for 55 minutes and further rinsing (3×30 minutes) in PBS with 0.3% Triton X-100 (PBT). Brains were then preincubated in 5% normal goat serum in PBT for 3 hours at room temperature with gentle agitation, followed by incubation in 1:50 dilution of biotinylated anti-lucifer yellow antibody (RRID: AB_2536191) in universal antibody dilution solution (Sigma Aldrich) for 3 days at 4°C with occasional gentle agitation. Brains were then rinsed (3×30 minutes) in PBT, followed by incubation with a 1:50 dilution of NeutraAvadin DyLight 633 for 3 days at 4°C. The samples were then rinsed in PBT, dehydrated through an ethanol series (70%, 90%, 100%, 100%), before clearing in methyl salicylate and mounting in a cavity slide using Permount.

Multiple overlapping Z-series of images covering the complete arborization of the injected neuron in both the brain and lobula complex were then obtained from the cleared whole-mount with a Zeiss LSM510 meta confocal microscope, using a 633 nm laser and 25× oil immersion objective (LD LCI Plan-Apochromat 25×/0.8 Imm Corr DIC; Zeiss) with a pixel resolution of 0.3×0.3 μm and optical sections every 1.5 μm in the 3^rd^ (Z) dimension. Overlapping image stacks were then stitched into a single volume using a plugin for ImageJ as described by Preibisch et al., (2009).

To better visualize the 3^rd^ dimension of the lobula dendritic tree (**Figure 1**), and the boundaries of synaptic neuropil, we post-processed the brain in order to counterstain the synaptic neuropils using an anti-synapsin antibody (RRID:AB_528479). The coverslip was removed and the brain dissolved out from the Permount by immersion in xylene (3 hours at room temperature). Following rehydration through a descending ethanol series and resuspension in PBT, the brain was then prepared for vibratome sectioning by embedding in a gelatin-albumin mixture (4.8% gelatin and 12% ovalbumin in water) which was allowed to set before post-fixing overnight in 4% Paraformaldehyde at 4°C. 200μm horizontal sections (i.e. at a right angle to the images shown in **Figure 1A-C**) were then cut on a Leica vibratome and rinsed in PBT. Sections were then blocked using 5% normal goat serum in PBT, before incubation with anti-synapsin for 3 days at 4°C in the dark. After rinsing with PBT (6×20 minutes) sections were incubated in secondary antibody (goat anti-mouse conjugated with CY3) at a dilution of 1:300, for 3 days at 4°C. The incubation solution also contained a 1:50 dilution of streptavidin conjugated CY5 in order to refresh fluorescence of the anti-lucifer antibody. Sections were then rinsed in PBT (6×20 minutes) before clearing and mounting in Rapiclear 1.49 with a 200μm spacer between the slide and coverslip (both from SUNJin Lab). 2× oversampled (in all 3 dimensions) image stacks from the ventral lobula arborization were then obtained using a Leica SP8 DLS confocal microscope in ‘Hyvolution’ mode, with 0.5 Airy unit pinhole and using a 20x oil immersion objective. These were subsequently deconvolved using Huygens Essentials software. Resulting stacks were then imported into Neutube 1.0 (Feng et al., 2015) for rotated volume rendering (2 representative rotations shown in **Figure 1D**).

### 1.2 Dendritic analysis

The imaged volume was imported into Neutube 1.0 (Feng et al., 2015) and dendrites were traced manually to reconstruct a 3-dimensional model for the main branches, bifurcations and terminals (Table 1). The reconstructed compartments were saved out in SWC format and subsequently rotated to a plane orthogonal to the relatively flat main arborization, using Matlab 2016B. A similar SWC model for HSE was downloaded from a publicly available database (Ascoli et al., 2007). The main dendritic (presumed input) arborization in the lobula / lobula plate of both neurons was then used for subsequent modelling. In NEURON, there are two types of geometrical structures, termed ‘segments’ and ‘sections’. A segment corresponds to an electrical compartment. Sections are unbranched and continuous lengths of cable consisting of a number of segments (Hines and Carnevale, 2001). The number of morphological (SWC) 3D points obtained were 4026 and 1697 for BSTMD1 and HSE respectively. The NEURON functions called Import3d_SWC_read and Import3d_GUI were used to convert continuous 3D point branches into sections which were then connected. This gave 692 sections for BSTMD1 and 576 for HSE. Each section was allocated one segment, i.e. a single electrical compartment. The projected center position of each section was then extracted and fed into the first (ESTMD) stage of the hybrid computational model, implemented in Matlab 2016B. Thus, each section has one synapse at this center position to receive input from ESTMD model.

**Table 1.**
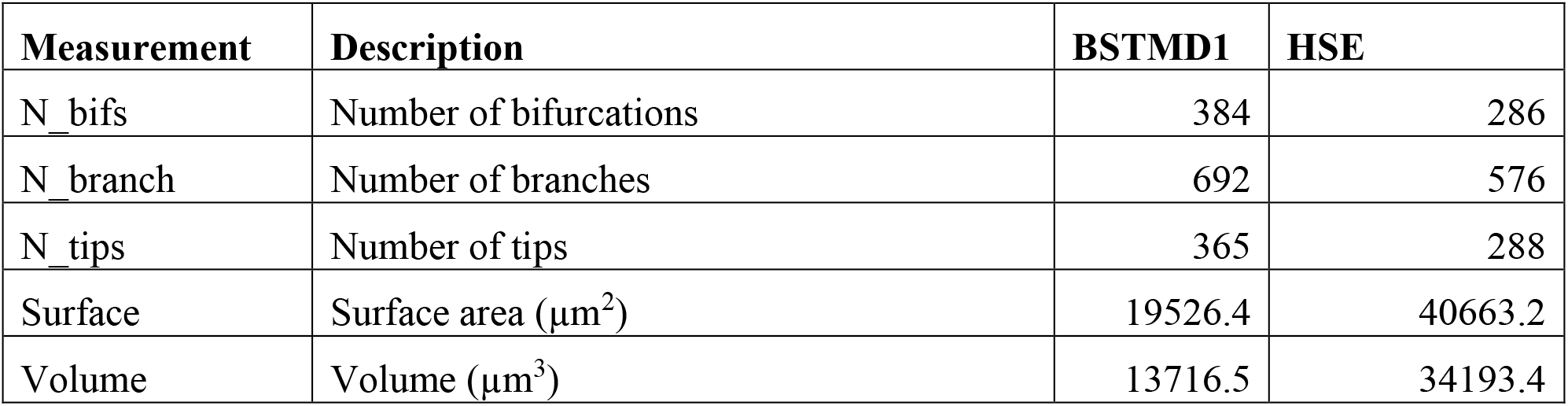
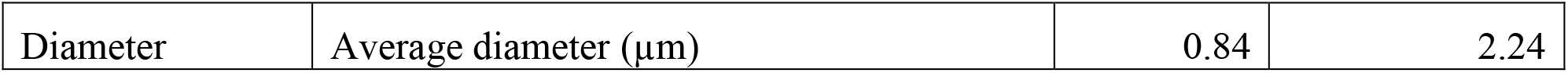
A selection of dendritic metrics for each dendritic tree.

| Measurement | Description | BSTMD1 | HSE |
| --- | --- | --- | --- |
| N_bifs | Number of bifurcations | 384 | 286 |
| N_branch | Number of branches | 692 | 576 |
| N_tips | Number of tips | 365 | 288 |
| Surface | Surface area ($\mu\text{m}^2$) | 19526.4 | 40663.2 |
| Volume | Volume ($\mu\text{m}^3$) | 13716.5 | 34193.4 |
| Diameter | Average diameter ($\mu\text{m}$) | 0.84 | 2.24 |

An alternative method would be to create one section with one segment for each 3D point and then connect them. This way one section would no longer correspond to one continuous branch but instead one 3D point. However, the number of continuous branches is less biased compared to the number of 3D points, since each 3D point is placed manually at arbitrary locations or using a local fitting algorithm when reconstructing the neuron. We thus used the continuous branches as sections rather than individual 3D points when placing input receptors. Another alternative method would be to create a number of segments in each section based on length of the branch. This way the exact 3D positions along a branch is lost but is roughly compensated for by addition of a branch length-based number of segments. One advantage of the method we selected is that the number of continuous branches are many fewer than the number of 3D points from the original neuron tracing, which enabled high computational efficiency.

### 1.3 Hybrid computational model approach

The overall architecture of our modelling approach is illustrated in **Figure 2**. The first stage is a bioinspired “elementary STMD” (ESTMD) model, which has previously been applied in robotics simulations for target tracking in visual clutter (Bagheri et al., 2017). The model was based on a parametric model previously shown to provide a quantitatively good match to the tuning properties of dragonfly STMD neurons (Wiederman et al., 2008). This model accounts for early visual processing by the photoreceptors and lamina cells, and then for target matched filtering by local small-field elements which are presumed to be an array of retinotopically organized neurons (‘ESTMDs’) that lie on the inputs to higher order STMD neurons (Wiederman et al., 2008). The outputs of this model then provide the input to a biophysically plausible multicompartment model for dendritic integration by a higher order neuron such as BSTMD1, implemented in the NEURON simulator (**Figure 2**).

**Figure 2.**
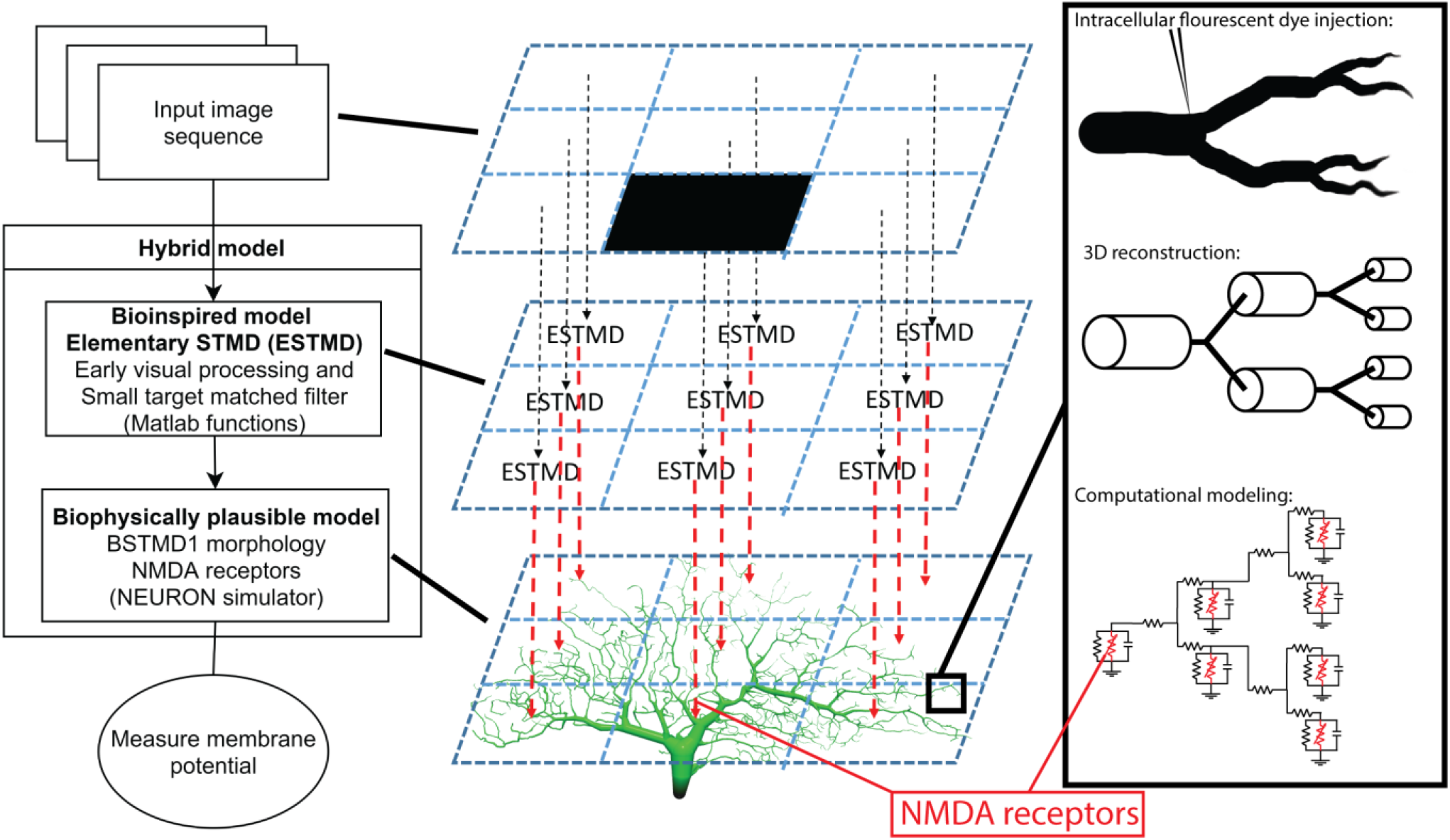
Illustration of the hybrid model. To the left is an overview of the model and what type of processing and mechanisms each component consists of. The middle column is a simplified explanation of how an input image is mapped on to the dendritic tree. To the right is a magnified illustration of the dendritic tree and how it was simulated, from dye injected neuron, reconstructed cable (cylinders) model, to mathematical compartment (electrical circuit) model.

#### 1.3.1 ESTMD model stage

The ESTMDs in our hybrid model are implemented as a set of mathematical functions in Matlab that take a matrix (any arbitrary luminance image sequence) as the input and generates an output image sequence. In the early visual processing box (**Figure 2**, left) the green channel from RGB input imagery is extracted, blurred and subsampled followed by temporal and spatial bandpass filters to mimic the photoreceptors and lamina of the insect optic lobe and reject redundancy from the image. The next stage of processing is presumed to represent columnar neurons within the medulla or the outer (primary) lobula (**Figure 1A**), since these lie just distal to the inputs or large and small field target-selective neurons in both dragonflies and flies (Nordström et al., 2006; Barnett et al., 2007; Wiederman et al., 2017). A matched filter for small-targets is then constructed as follows: First, we separate the response to brightening events (‘ON’) and dimming events (‘OFF’) into different parallel pathways by half-wave rectification of the input signal (with inversion of the negative phases). In the ESTMD model, movement of a small contrasting feature is assumed to consist of a stimulus that triggers either an ON or OFF detector in one half of the pathway (the leading edge of the feature) followed by an opposite sign stimulus with a short delay at the same location as the trailing edge passes the same location in space. A low-pass temporal filter delays the signals from each partially rectified detector so that a non-linear (multiplicative) correlator within the ESTMD compares each delayed signal with the undelayed signal of opposite sign. Combination of this ‘feature template’ with fast adaptation (to reject background texture) and center-surround antagonism provides a sharp selectivity for small, moving targets within the input images (Wiederman et al., 2008; Bekkouche et al., 2017). The output of this model stage was then mapped onto the presumed input dendrites of the neurons, as projected into a 2-dimensional image (**Figure 1B**) assumed to be a retinotopic projection of the space in the visual field.

#### 1.3.2 Connectivity and stimulus sequences

The ESTMD model is fed with a sequence of images containing an animation of a translating black target of 30×30 pixels within a 960×540 pixel field which was mapped onto the neuron dendritic tree. Assuming that the input field corresponds to 60×60 ° (ESTMD model setting) of the visual field of the neuron (Dunbier et al., 2012), this corresponds to a target of angular size 1.88×3.33 °, a size which is close to that determined by several studies to be a near-optimal stimulus for dragonfly STMDs studied *in vivo* (O’Carroll, 1993; Wiederman et al., 2017; Fabian et al., 2019). The early stages of image processing within the model blur the image to represent blur in the optics of the insect eye, and the images are subsequently subsampled by the ESTMD model to a matrix of 34×60 pixels, corresponding to sampling by the compound eye (Bagheri et al., 2015, 2017). The ESTMD algorithm then represents the detection of any moving targets through increased values in the output target matrix. Since insect small field STMD neurons typically exhibit very low (or no) spontaneous activity in the absence of moving targets (O’Carroll, 1993; Barnett et al., 2007; Wiederman et al., 2017), we then simulate a spike threshold at this stage by continuously filtering the target matrix with a threshold of approximately 25-50% of the typical maximal ESTMD output when a target was present somewhere in the scene. This was empirically determined for this type of stimulus (moving black targets), such that only the ESTMDs near a target generate responses above this threshold. The threshold enables the model to ignore weak early target motion responses at levels which could be generated by for example random flickering targets or a moving bar. The maximum value among these filtered positions is then selected as an estimation of the target position on each frame. The estimated position is then used to give each synapse a probability of receiving an input impulse, based on the normalized distance from the synapse to the estimated target location according to the following function:

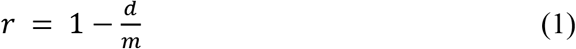

Where d is the distance from a receptor to the target, m is the maximum distance and is set equal to the diagonal of the subsampled image. The r is thus a normalized inverse distance between 0 and 1 and corresponds to the probability of an input spike decreasing linearly with the distance. This value is then inserted to a Gaussian curve function (Gaussmf function in Matlab 2017a) that performs the following conversion from linear to Gaussian probability:

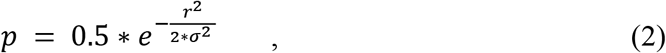

where 0.5 is a tuning variable that was manually set to generate a maximum of around 100-300 input spikes per synapse and *σ* = 0.1 generating ESTMD neighbor overlap at half max of 14.1 °. A uniformly distributed random number generator function called rand (rand in Matlab 2017a), generating numbers between 0 and 1, is then used to determine whether a certain synapse should receive an input spike or not:

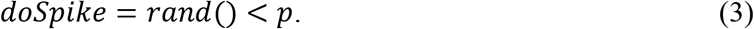

#### 1.3.3 BSTMD1 (dendritic integration) model stage

The second part of the model is biologically plausible and consists of the anatomical model for the dendritic tree of the BSTMD1 neuron (Figure 1B), implemented as a multicompartment model in the NEURON simulator.

The NEURON compartmental model is based on the following equation:

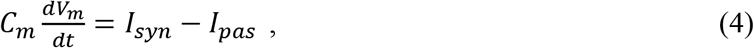

where the membrane capacitance Cm =1F/cm^2^ and *I*_*pas*_ is the passive membrane leakage current:

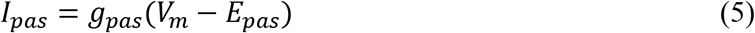

Synaptic current at time *t* after activation is given by:

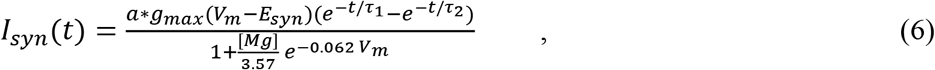

where synaptic reversal potential *E*_*syn*_ = 0, magnesium concentration [Mg] = 1mM, normalized maximum conductance *g*_max_ = 1, rising time constant *τ*_1_ = 4*ms*, falling time constant *τ*_2_ = 42*m*. The variable a is chosen so that the maximum value of the synaptic conductance *I*_syn_/(*V*_*m*_ − *E*_syn_) matches *g*_max_. More information about this synaptic model can be found in Baker et al. (2011). The characteristics of the NMDA synapse are illustrated in **Figure 3A** and **B**. The passive membrane variable values were taken from Shoemaker (2011) and the default NMDA variable values were used unless otherwise stated.

**Figure 3.**
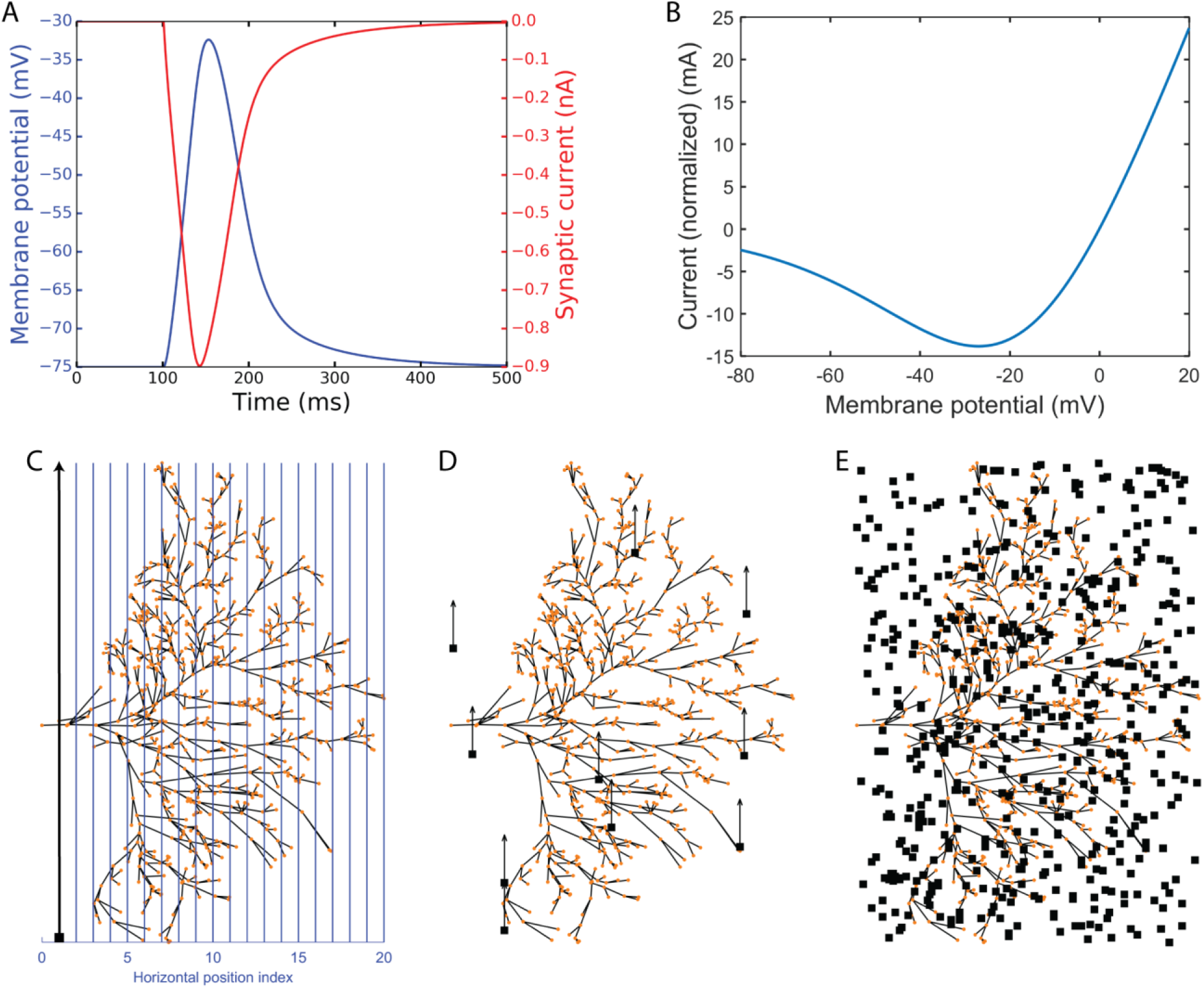
Experimental simulation protocols. **(A,B)** illustrates NMDA synapse characteristics. **(A)** Impulse response showing membrane potential (blue) and synaptic current (red). **(B)** Plot of the macroscopic NMDAR current (shown as normalized current). It is essentially all the voltage dependent parts normalized by multiplying by 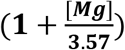. **(C-E)**. Illustration of the experimental protocols. The red dots represent the dendritic sections which have been given one NMDA synapse each. The neuron viewed from the anterior side of the brain (**Figure 1A**). **(C)** The continuous experimental setup where blue lines show the vertical trajectories on which the small target (black target) travels (in the direction of the black arrow) with 500 steps (ms). **(D)** The short experimental setup where a target starts at a random position and travels up for 50 ms, switches to a new random position, travels up for 50 ms again and repeats this 10 times in total (500 ms). **(E)** The random experimental setup showing an example of 500 random positions (500ms).

To illustrate the membrane potential dependence of the macroscopic NMDAR current the equation 7 was extracted using the voltage-dependent parts of equation 6, and then normalized by multiplying by 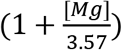 so that when *V_m_* = 0, *I_norm_* = −*g_max_ E_syn_* = 0 (because *E_syn_* = 0). The result, plotted in **Figure 3B** is the following equation:

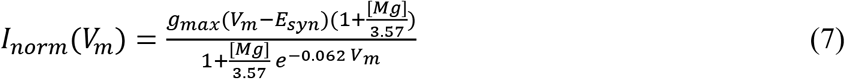

This normalization sets 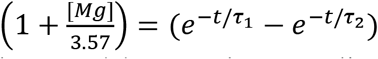 thus showing synaptic current at the specific 3.57 time (and tau’s) in which *I*_*norm*_(0) = 0. The normalization does not restrict values between two values (for example 0 and 1). Thus, we can use the unit mA as shown in **Figure 3B**.

For most simulations, the final output of the model was the graded membrane potential, *V_m_*, estimated for the most basal section of the multi-compartment model (i.e. the location indicated by the ‘x’ in **Figure 1B**). This represents the integrated signal of the main dendritic tree, and thus is also the generator potential for spike generation.

#### 1.3.4 Spiking model output

Because BSTMD1 is a spiking neuron, we also developed a spiking model variant to allow us to compare model outputs with spike-trains recorded from BSTMD1. Active sodium (*I*_*Na*_) and potassium (*I*_*K*_) ion channels, based on a default Hodgkin Huxley model mechanisms (Hodgkin and Huxley, 1952) from a software application called Neuroconstruct (Gleeson et al., 2007), were used in the experiments with spiking properties. The model in those experiments can be described by the following equations:

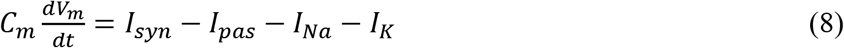

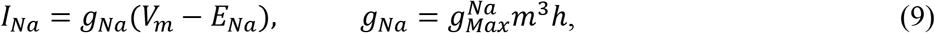

where *E_Na_* = 50 *mV*, 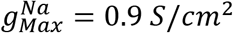

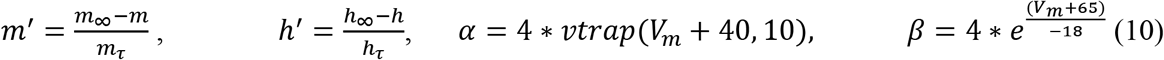

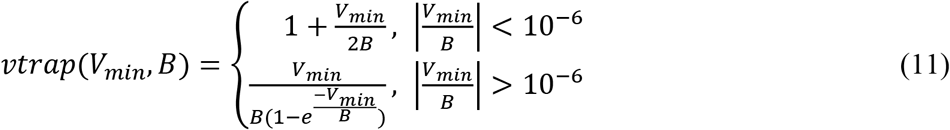

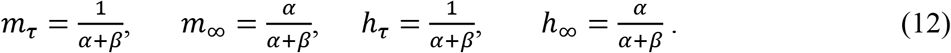

Similarly, for potassium

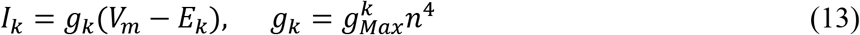

where *E*_*k*_ = −85 *mV*, 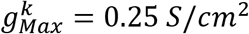

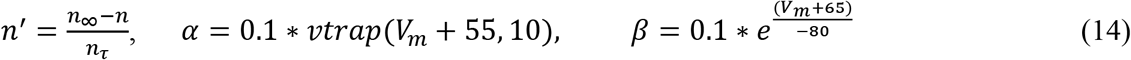

In the spiking version of the model, one sodium and one potassium mechanism were placed on each section where the relative diameter > 90% of the maximum diameter. This criterion resulted in 6 sections for the BSTMD1 model and 5 for HSE.

## 2 Results

### 2.1 Responses to targets moved on continuous or discontinuous paths

We first tested whether our model is able to capture the facilitation observed in dragonfly STMDs – a boost in the local response gain when targets are moved along long continuous trajectories. We compared responses to such trajectories with shorter path stimuli distributed across the dendritic tree (i.e. discontinuous paths). Our stimulus protocol is illustrated in **Figure 3C-E** and the supplementary video S1. In the ‘continuous’ path condition, the target moved from the bottom to the top of the image, corresponding to a presumed ventral to dorsal trajectory for targets in the real world (the specific retinotopic mapping to the inner lobula neuropil where BSTMD1 arborises remains to be established). Each trajectory comprised 500 images, animated at 1000 frames per second, hence eliciting 500ms of target motion from the bottom of the dendritic tree to the top at its broadest point. A sequence of 20 such paths was tested, shifted horizontally across the dendritic span, presumed to map to different horizontal positions (e.g. from anterior to posterior, **Figure 3C**). The only stochastically varying factor was the Gaussian distributed synaptic input, generated from the ESTMD model output as input to the NMDA synapses on the BSTMD1 or HSE neuron. Nevertheless, each of the 20 paths was replicated 3 times, to check that the stochasticity of the model does not introduce large variability into the output for a given path. In the ‘short’ path condition, targets moved on 10 shorter trajectories, each lasting 50ms, before the target was displaced to a new location within the visual field at random, again for a total of 60 trials each containing 500ms of such discontinuous target motion (**Figure 3D**). In the ‘random’ condition, which provided a control for the model rejecting uncorrelated target flicker, the target was displaced to a new random location on each video frame, again repeated 60 times (**Figure 3E**). We also executed additional conditions, representing intermediates between the short path and fully random stimuli, using paths of 5 or 25ms duration.

**Figure 4** shows simulations of neuronal activity for our hybrid models for both BSMTD1 and HSE in response to these 3 stimulus conditions. As expected, the random (flicker) target stimulus produced no measurable change in membrane potential at the generator location (identified by an ‘x’ in **Figure 1B**) for any model variant, seen as black lines of constant resting membrane potential in the left column of **Figure 4**. In our initial modelling of NMDA synapses with constant synaptic weight (8.25 pS) we found that different variants of the model gave different output levels. The strongest outputs were seen for the BSTMD1 model (**Figure 4A**), while the HSE model output was much weaker (**Figure 4B**). In BSTMD1, individual continuous paths produced the strongest responses, and these build over time, reaching a peak towards the end of the 500ms stimulus period (**Figure 4A**, blue and orange lines) and then fading away as the continues to move beyond the ‘receptive field’ defined by the dendritic structure. Stimuli that transected the central part of the dendritic tree (e.g. positions 6-12 in **Figure 3C**) gave stronger and more sustained responses leading to both higher peak and mean depolarization (**Figure 4A**, right). Shorter (50ms) stimulus paths produced responses that often peaked earlier and then plateaued at a lower final level (**Figure 4A**, red lines). Average values were calculated from the gray area in **Figure 4** left column. Boxplot analysis (**Figure 4** middle column) using these average values for all trajectories shows a higher median and large interquartile range for the long path data versus short paths. We segregated a subset of continuous paths from the receptive field center, based on a criterion of depolarization to at least 50% of the peak response seen (**Figure 4** orange). Analysis of this subset of long paths confirms the large facilitation effect, with continuous paths giving significantly higher average responses than short path trials (*p* = 1.12 ∗ 10^−11^, rank sum test, **Figure 4A**, middle).

**Figure 4.**
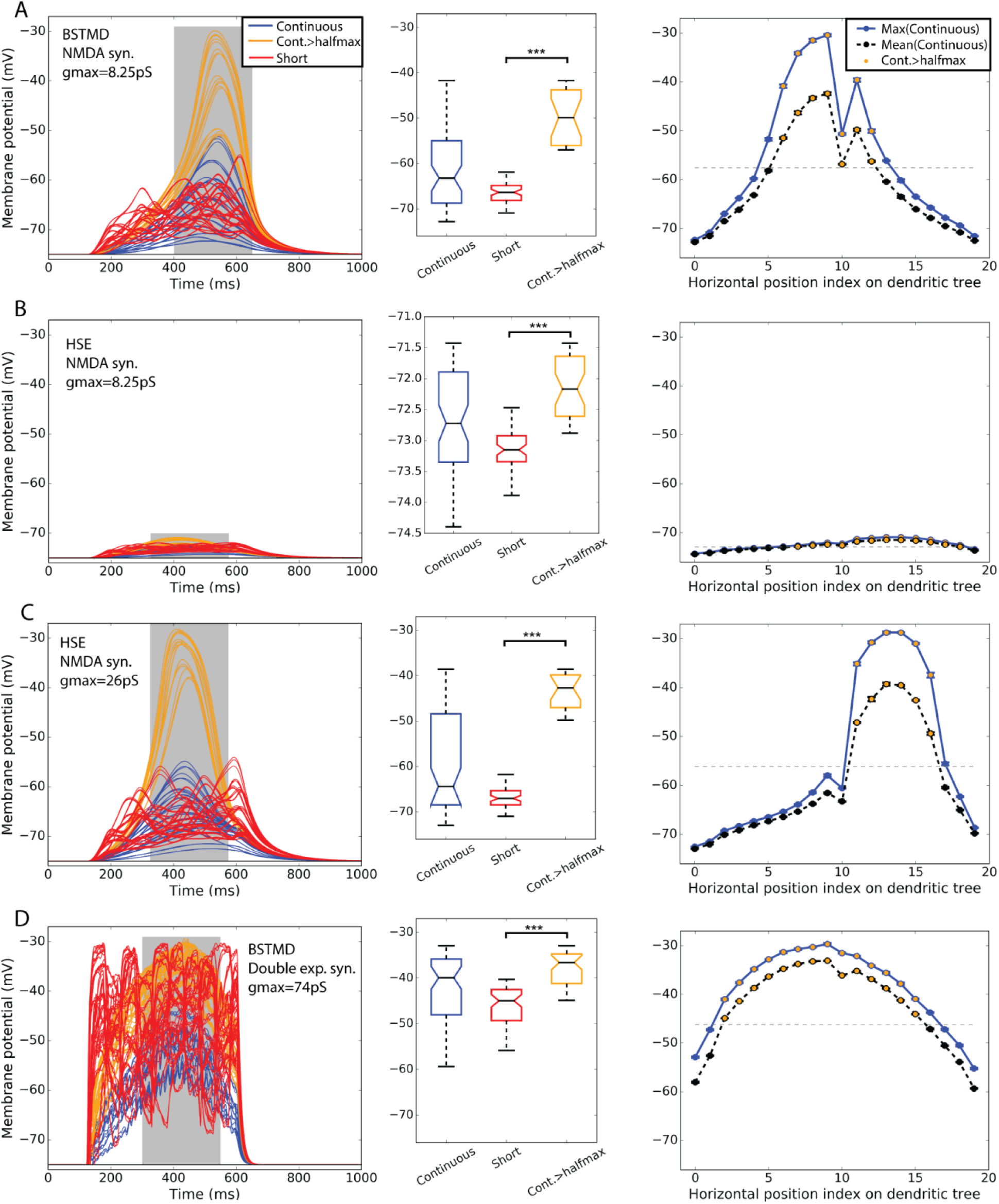
BSTMD1 and HSE simulations indicate difference in bimodality. Row **(A)** shows BSTMD1 simulations. **(B and C)** shows HSE simulations, one with the same synaptic gain (Gmax) as the BSTMD1 simulation and one with higher synaptic gain. **(D)** shows BSTMD1 simulations with double exponential (exp2syn) synapses instead of NMDA synapses. The left column shows the membrane potential over time. The middle column shows a box plot of the average membrane potentials of the curve area marked in gray in the left column. The right column shows average maximum (entire curve) and average (from gray area) membrane potentials (and SEM) of three identical trials differing only in randomness from spike input generation. There was not much variance between the trials so the SEM near the data point marker. ***A: ***p* = 1.12 * 10^−11^ B: *p* = 8.52 * 10^−14^, C: *p* = 1.52 * 10^−10^, D: *p* = 8.08 * 10^−13^**. The median difference for Cont.>halfmax vs. short: A=16.49, B=0.98, C=24.35, D=8.37 (mV).

At position 10 for the continuous path experiment, responses show a conspicuous dip compared with the adjacent paths, seen across all models (**Figure 4**, right column). This results from a mismatch (aliasing) between the alignment of target trajectories in the input images and the inputs of the ESTMD model. This aliasing reduces the number of animated frames in which target features exceeded the threshold required to be detected by the ESTMD model, resulting in a locally weaker response. Although we could have re-run the models with more perfectly aligned trajectories to eliminate this artefact, the presence of this less strongly activated path, well within the main anatomical receptive field, revealed important features of the local spread of facilitation which we will describe below (section 2.5).

In the HSE model, we observed much weaker responses for the same gain as the BSTMD1 dendritic tree (**Figure 4B**). Continuous path stimuli sometimes produced even weaker responses than typical short-path trials, but these were mostly from stimulus paths that moved across the parts of the dendritic tree with lower synaptic density (paths 0-10 and 17-19, **Figure 4B** right column). Paths that transected larger patches of input dendrites (paths 11-16) produced stronger facilitation than the short path stimuli, although the additional depolarization is only on the order of 1mV, compared with >15mV seen for continuous paths in the BSTMD1 model. To be sure that the reduced facilitation in the HSE model was not due to the weaker overall activation, we tested an additional HSE model variant where we increased the synaptic gain by 3.15x (to a conductance of 26pS per synapse, versus 8.25pS in the BSTMD1 model). This allowed the HSE model to reach similar peak membrane potential levels to BSTMD1 after several hundred milliseconds of target motion. This also led to an increase in the facilitation effect, with the continuous path responses forming two clusters, one more strongly facilitated than the other. Again selecting a subset of trials corresponding to the effective receptive field center (responses > half maximum, orange lines) we observed significant facilitation compared with the short path trials in both the low and high gain variants of the HSE model (**Figure 4B**, *p* = 8.52 ∗ 10^−14^, **Figure 4C**, *p* = 1.52 ∗ 10^−10^, rank sum test).

### 2.2 NMDA receptors are necessary (but not sufficient) for facilitation

To confirm that NMDA receptors are required to elicit the facilitation effect in BSTMD1, and that it was not due to some other morphologically dependent process of integration, we also tested a variant of the BSTMD1 model that employed a classic form of a double exponential synapse, instead of NMDA receptors (**Figure 4D**). The main difference is that the NMDA synapse have the magnesium block function and longer rise and decay times. We needed to increase the synaptic gain of the double exponential synapse model almost 9-fold (to 74mS) compared with the NMDA version, in order to reach similar levels of depolarization. We still observed weaker facilitation in the time course for this model for any trajectory (**Figure 4D**). A subset of the continuous trials from the receptive field center still gave significantly higher responses than the short path trials (**Figure 4D**, *p* = 8.08 ∗ 10^−13^, rank sum test). However, even during the short segment trials, most stimulus paths include segments that correspond to more dense parts of the dendritic tree and despite their very short duration, these still elicit transient depolarizations to similar peak levels as those seen during continuous trajectories in the majority of such trials (**Figure 4D** left, red lines). This suggests that NMDA receptors are necessary to elicit the clear time-dependency in response build up seen in both the BSTMD1 and HSE model variants. Since such slow buildups were never observed in the NMDA models for shorter paths, we further conclude that NMDA receptors alone are not in themselves sufficient to produce this facilitation. Rather it requires the sequential activation of such receptors on nearby dendrites for continuous paths. One important point here is that the HSE model has just 16.8% fewer synapses compared to the BSTMD1 model, yet required a more than 3-fold increase in synaptic gain to generate the same membrane potential. This suggests that the BSTMD1 morphology may be better optimized for continuous object tracking than HSE.

### 2.3 Dendritic morphology analysis

Does the weaker facilitation in HSE compared with BSTMD1 result from differences in the spatiotemporal synaptic integration, i.e. the interaction between the nonlinearity of the NMDA synapses and the specific morphology of the dendritic tree structure in BSTMD1? The total spatial field occupied by HSE dendrites is larger than BSTMD1, and is more spread out along the medial-lateral axis (**Figure 1B**), reflecting differences in the projection from its inputs (in the primary lobula and medulla of the fly) to the retinotopic map of the integrating dendrites within the dipteran fly lobula plate, where HSE arborizes (Hausen, 1982). While the outer (primary) lobula of the dragonfly is substantially larger than its dipteran fly counterpart, spreading across more than 1000μm in its ventral to dorsal extent, BSTMD1 sits within a deeper neuropil, the medial lobula (**Figure 1A**). It has a much more compact and very dense dendritic tree, with a locally much higher density of dendrites than HSE (**Figure 1**). Despite its smaller physical extent, this greater dendritic density leads to our BSTMD1 model still having more total sections (and thus compartments in our simplified model) than HSE (692 versus 576). This in turn leads to a larger number of NMDA receptors in the BSTMD1 model, since we assumed 1 receptor/section in the modelling protocol. That in itself likely creates some differences regarding input-response balance, although if we assume that the space constant is at least similar to the physical length of each section, this can easily be compensated by adjusting the input weights, as we did for the second HSE model variant. The dendritic length difference seen between BSTMD1 and HSE in **Figure 1B** (along the medial-lateral axis) should not matter significantly, since the current modelling protocol scales the stimuli to span across the extent of both dendritic trees.

To further quantify the morphological differences between these dendritic trees extracted metrics from our anatomical models of the dendritic trees. Further explanation of these metrics can be found in Scorcioni et al. (2008) (Lmv5.3 software help folder). The main results (**Table 1**) confirm that the number of bifurcations, branches and tips are all lower in HSE despite its larger overall size. A larger number of branches affects the distribution of the inputs and therefore limits the precision with which a target can be tracked. The surface area and volume in HSE is also more than double that in BSTMD1 (**Table 1**), meaning that input currents need to spread further to affect neighboring NMDA receptors. The average diameter of neurites is also much lower in BSTMD1 (Table 1). Diameter is proportional to the length constant and (and thus also to conduction velocity for actively propagating signals) due to reduced axial resistance (Pumphrey and Young, 1938). This would enhance nonlinear interaction between synapses in the HSE neuron versus BSTMD1, yet we see the opposite in terms of facilitation, suggesting that the much higher density of synapses that are activated sequentially in BSTMD1 more than compensates for the reduced length constant.

From the above analysis we conclude that the large difference in recruitment of facilitation between these neuron types indeed results from the unique and very compact and dense dendritic arborization in BSTMD1. In this context it is worth noting that fly lobula plate HS neurons (including HSE) are primarily graded neurons, which conduct excitatory or inhibitory membrane potentials (up to +/− 20mV) right to their axon terminals (Hausen, 1982). This requires large diameter neurites, including those extending to the inputs right at the periphery of the lobula plate. These are well within the resolution of optical microscopy in fixed tissue and in some cases even for live imaging of dendritic calcium signaling (Single and Borst, 1998; Dürr and Egelhaaf, 1999; Haag et al., 2004). Hence anatomical models for these neurons are relatively complete. By contrast, BSTMD1 is primarily a spiking neuron (Dunbier et al., 2012). Although its main axon is also very large, BSTMD1’s very compact main dendritic tree includes numerous very fine neurites which remain a challenge to image well in whole mount preparations (**Figure 1C**), even when applying state of the art tissue clearing and confocal imaging techniques (Bekkouche et al., 2020). We were thus not able to accurately trace every fine neurite. Furthermore, in the process of translating our anatomical model into the Neuron simulator based on those dendrites that were traceable, it was also necessarily to down-sample the number of segments. Consequently, if anything our models underestimate the difference between these two neuron morphologies and their effect on nonlinear interactions. Hence an even more realistic model for the complete BSTMD1 tree would be expected to show even stronger facilitation than our current model suggests.

### 2.4 Priming of facilitation and comparison with BSTMD1 intracellular responses

Priming stimuli have recently been used in several studies to link neuronal facilitation to neural mechanisms of selective attention (Wiederman et al., 2017; Lancer et al., 2019). When a target is moved initially along a long trajectory (a ‘primer’) and then displaced to new locations within the receptive field to map the local change in response compared to short-segment (‘probe’) stimuli, the neuron shows enhanced sensitivity to the probe stimuli selectively at locations close to the last seen location, while responses are suppressed elsewhere (Wiederman et al., 2017). To compare intracellularly recorded neuronal responses with the BSTMD1 model, we reproduced this experiment for the spiking variant of our hybrid model (**Figure 5). Figure 5A, B** shows the response of the recorded neuron and the BSTMD1 model to either the probe stimulus alone (‘Probe’, blue line) or to probe stimuli preceded by a 500ms primer (‘Primer + Probe’, red line). The same data are plotted here both as raw responses (upper), and as instantaneous spike frequency plots (1/inter-spike interval, lower). Even though the probe stimulus was placed within a highly sensitive part of the receptive field (as indicated by the vigorous response to the primed stimulus by the time the target reaches the probe location) the response to the probe alone shows the characteristic slow rise in response over several hundred milliseconds before the spike frequency begins to quantitatively resemble the facilitated (Primer + Probe) condition. This facilitation onset is also evident at the commencement of the primers, for both the neuron and BSMTD1 model. This shows that the neuronal facilitation time course is well matched by our NMDA-based model for BSTMD1. Note that in this case, we needed to increase the gain of the BSTMD1 synapses to a higher value than in our earlier simulations (40pS), in order to achieve similar spike rates to those observed in the neuron. Other means of adjusting the spike rate is by changing the neuronal or synaptic properties such as the conductance and reversal potential of the voltage-gated sodium and potassium channels, membrane resistance, capacitance and synaptic temporal constants. Such adjustments can also have other effects on the neuronal excitability and the synaptic transfer function leading to irregular firing dissimilar from what was observed in the BSTMD1. The parameters were manually tuned to mimic the BSTMD1 spiking, limiting the ranges in which each parameter could be used to adjust for spike rate.

**Figure 5.**
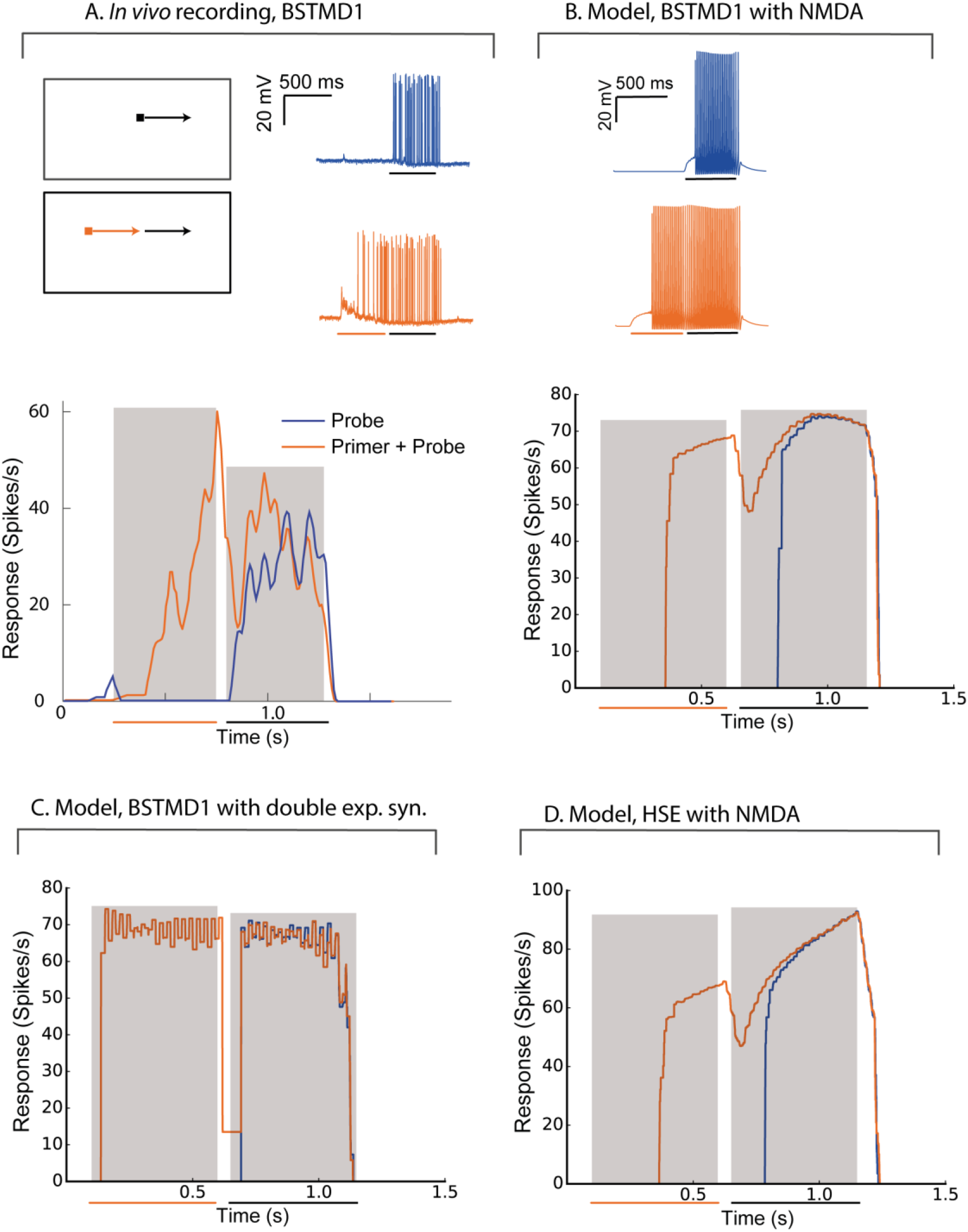
The model behaves similar to intracellular recordings of BSTMD1. Example BSTMD1 intracellular electrophysiology data of stimuli involving a probe (moving target) with or without a primer (moving target before the probe appears). **(A)***In vivo* BSTMD1 recording of primer+probe experiment. **(B)** BSTMD1 model (gmax=40pS) with NMDA synapses. **(C)** BSTMD1 model (gmax=120pS) double exponential synapses instead of NMDA synapses. **(D)** HSE model (gmax=135pS) with NMDA synapses.

Part of the long onset latency (i.e. before the first spike is seen) may be due in part to the threshold that was introduced into our model to reduce the response in the absence of target stimuli. This threshold was necessary to mimic the very low spontaneous activity evident in the neuron data during the pre-stimulus period (**Figure 5A**). This together with the slow kinetics of the NMDA receptors seem to have delayed the onset of the response by around 150-250ms. When we used a similar spiking hybrid model, but now applied to the output of the double exponential synapse model (with gain 120pS), the facilitating time course is no longer evident (**Figure 5C**), again confirming that the NMDA receptors are necessary. The absolute response latency is also shorter in this case. This is likely because the double exponential synapses require relatively high input weight to generate similar membrane potential response as when NMDA is used. If we use the high-gain HSE model (135pS), there is also a lag in response rise to the probe-alone condition, intermediate between the double-exponential and NMDA variants of the BSMTD1 models (**Figure 5D**). The high gain led to higher spiking during the end of the Primer + Probe condition and was necessary to generate spiking response of around 60-100 spikes/s during the priming period.

### 2.5 Spatial extent of the facilitation field

While our priming experiment reveals a similar time-course for response onset and facilitation to that observed in the BSTMD1 neuron, our model contained no specific attempt to localize the facilitation field to an area close to where the target was last seen, as has been observed in large field STMD neurons in the dragonfly (Wiederman et al., 2017). It is nevertheless possible that the limited signal spread between local compartments still produces a localized facilitation effect. To test whether this was the case, we estimated the extent of the facilitation field by locking the primer to a single path, at a horizontal position index of 10, corresponding to near the middle of the dendritic tree (see **Figure 3C**). Rather than traversing the entire dendritic field, we shortened the primer path to 50% of the previous vertical trajectory but kept the duration to 500 ms by reducing the target speed to half that as used in **Figure 4**, so that the primer path always ended in the middle of the receptive field (**Figure 6A, middle**). Following the primer, the target was then displaced to a Probe location (100ms duration, i.e. 10% of the vertical path) which was then varied in position systematically over a grid of 10 vertical locations for each of the 20 trajectories covering the entire simulated sensory input as previously illustrated in **Figure 3C**. One such trajectory is illustrated in **Figure 6A**. The response for each grid position is given as the maximum membrane potential from that period. Compared with prior studies on dragonfly neurons, this corresponds to the spike rate during a short period near to the onset of the probe response (Wiederman et al., 2017). For this study, the maximum-value was chosen since it enabled slightly better visualization contrast for the facilitation than the average membrane potential during the probe. We repeated this experiment for the same (non-spiking) BSTMD1 as in **Figure 4** (synaptic weight 8pS), as well as for an additional variant with increased synaptic weights (10 pS respectively) designed to better reveal the underlying ‘receptive field’ of the neuron in response to short probe stimuli presented with no primer, which otherwise elicited very small changes in membrane potential (**Figure 6**, left column).

**Figure 6.**
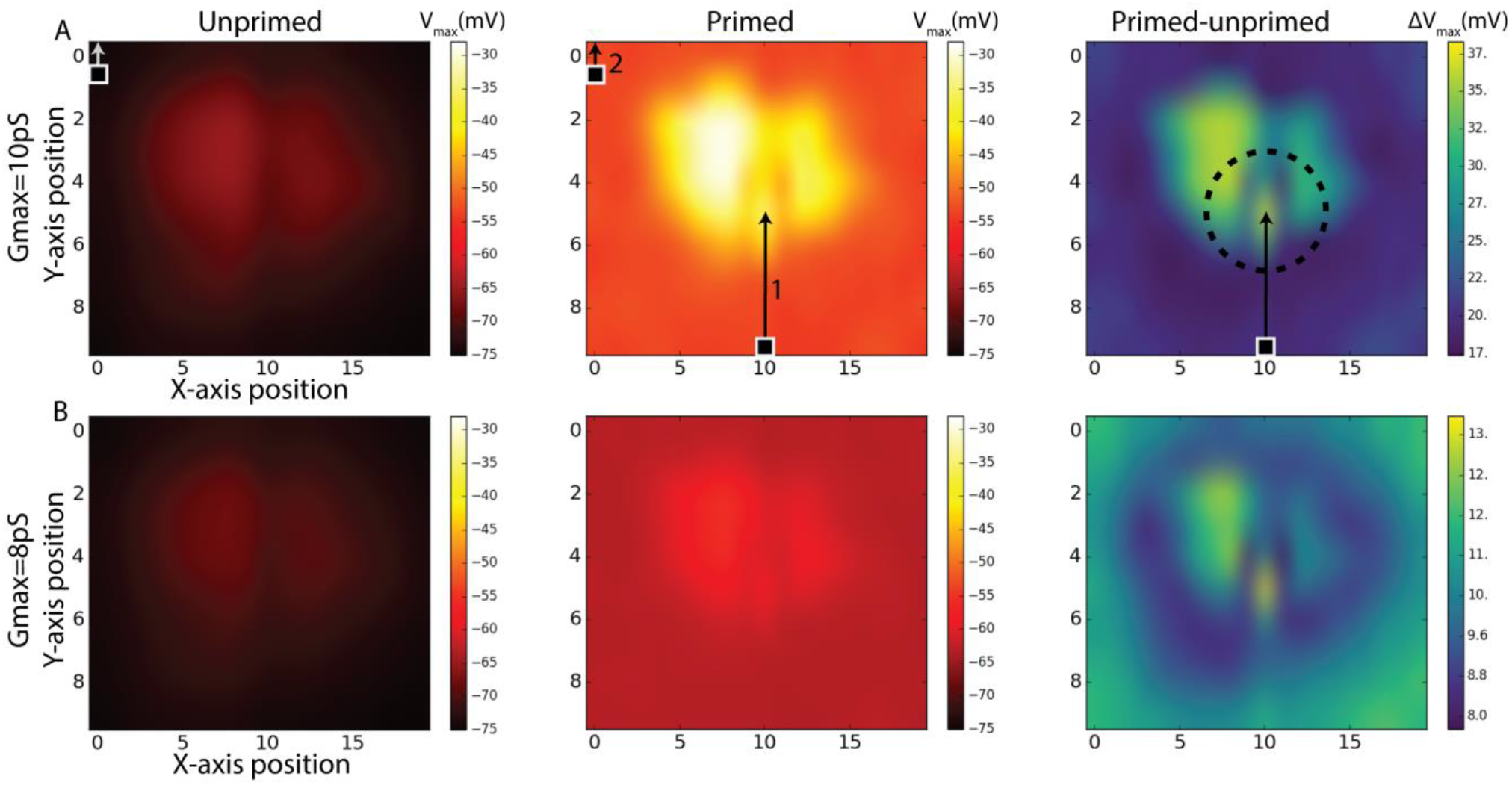
Simulations of Unprimed, Primed stimulus and the difference between the two (Primed-Unprimed). The response fields are mapped to correspond a view from the anterior side of the brain (**Figure 1A**). The rows **(A-B)** indicate the same experiment for different NMDA synaptic strengths (Gmax). The 100ms measurement period is indicated with a target square (probe) and arrow. In the first column (Unprimed) the probe position was varied systematically for each axis (10×20) position. The probe travelled for 100 ms corresponding to 10% of the entire vertical path. In the middle column (Primed) a primer (indicated by 1) preceded the probe (indicated by 2). Responses correspond to the maximum membrane potential in the non-spiking version of the model using the BSTMD1 morphology. The dashed circle in the right column (Primed-Unprimed) on row **(A)** indicate the area which could be expected to have higher activity (facilitation) due to the recent disappearance of the primer. This expectation can be expected for row **(B)** as well.

The middle panels in **Figure 6** show the effect on the underlying ‘receptive field map’ of adding the primer (path indicated by arrow 1 in **Figure 6A**). We further quantified this effect by computing the difference between the local responses in the primed and unprimed cases (**Figure 6**, right column) to estimate the facilitation field of the model (Right column, ‘Primed-unprimed’ **Figure 6**). This reveals that the response in the model is dependent on the local synaptic density. In addition, however, the primer induces a distinctive small ‘spotlight’ of locally enhanced activity near the terminal position of the primer, similar to that observed for very similar stimuli presented to dragonfly STMD neurons (Wiederman et al., 2017). Note that due to the previously mentioned aliasing/mismatch between input images and ESTMD model (section 2.1), the particular path that we selected for the primer actually produces weaker responses than adjacent paths, as clearly evident from the dark vertical band in the probe-only data (**Figure 6A**, left). Yet despite this being an intrinsically less sensitive path, the difference between primed and unprimed responses at this location is close to the maximum seen at any location (**Figure 6B**, right) thus confirming a highly localized spread of potent facilitation. In contrast to the specific location of the spotlight seen in the dragonfly data, which tends to lie slightly ahead of the target, this locally enhanced region of sensitivity is centered on the last seen location of the target in our data (indicated by the circle in **Figure 6A**, right). This spotlight is thus not ‘predictive’ of the future trajectory of the target, which is a key property of the reported dragonfly STMD facilitation field. We conclude that while dendritic spread of NMDA-mediated enhancement could potentially explain the *limited* spatial spread of the facilitation field observed in dragonfly neurons, this is not sufficient to explain the predictive nature of the latter.

### 2.6 The effect of direction on the facilitated response

To further investigate how the specific placement of NMDA receptors on the dendritic tree and the kinetics of facilitation interact for targets moving in different directions, we further tested the responses for facilitation across all 20 paths spread out over the horizontal input space as illustrated in **Figure 3C**. In each case, the stimulus moved either from bottom to top of this field (**Figure 7A-C**), or from top to bottom (**Figure 7D-F**). Stimuli either moved along the entire vertical trajectory over 500ms (long paths, **Figure 7B, E**), or were presented in one of 5 shorter (100ms segments) from the same paths (**Figure 7A, D**). Long upward paths produce maximal excitation (depolarization) further up in the stimulus space than for downward paths. To disambiguate which component of this response is due to the time course of facilitation, and which to the underlying receptive field structure, we then subdivided each of the responses from the long continuous paths into 5 shorter (100ms) periods and subtracted from this the corresponding short path response (**Figure 7C,F)**. Although the response to short path stimuli is weak for either direction (**Figure 7A, D**), remapping the color map for the upward direction stimuli (**Figure 7G**) reveals the intrinsic local sensitivity of the model, similar to the unprimed ‘receptive field’ in **Figure 6**. For comparison, we also show a map for the actual local synaptic density as extracted from our anatomical model, plotted onto the same 20×5 grid (**Figure 7H).** While the kinetics of the response introduced some temporal blur into this spatial pattern, the major features of this dendritic tree are still visible in the response field, with a broad area of strongest excitation corresponding the highest synaptic densities. This correlation is supported by a scatter plot for all 100 locations plotted, revealing a strong correlation (r=0.77, r^2^=0.60). Hence the difference plots (**Figure 7 C, F**) should account reasonably well for which component of local response enhancement is due to the underlying receptive field structure, and which is due to the build-up of facilitation. This clearly shows that the offset due to the stimulus direction results from the slow build-up of facilitation, rather than to any intrinsic sluggishness due to the kinetics of the underlying motion detectors. A similar shift in the apparent receptive field of the neuron due to facilitation has observed for the dragonfly CSTMD1 neuron (Nordström et al., 2011).

**Figure 7.**
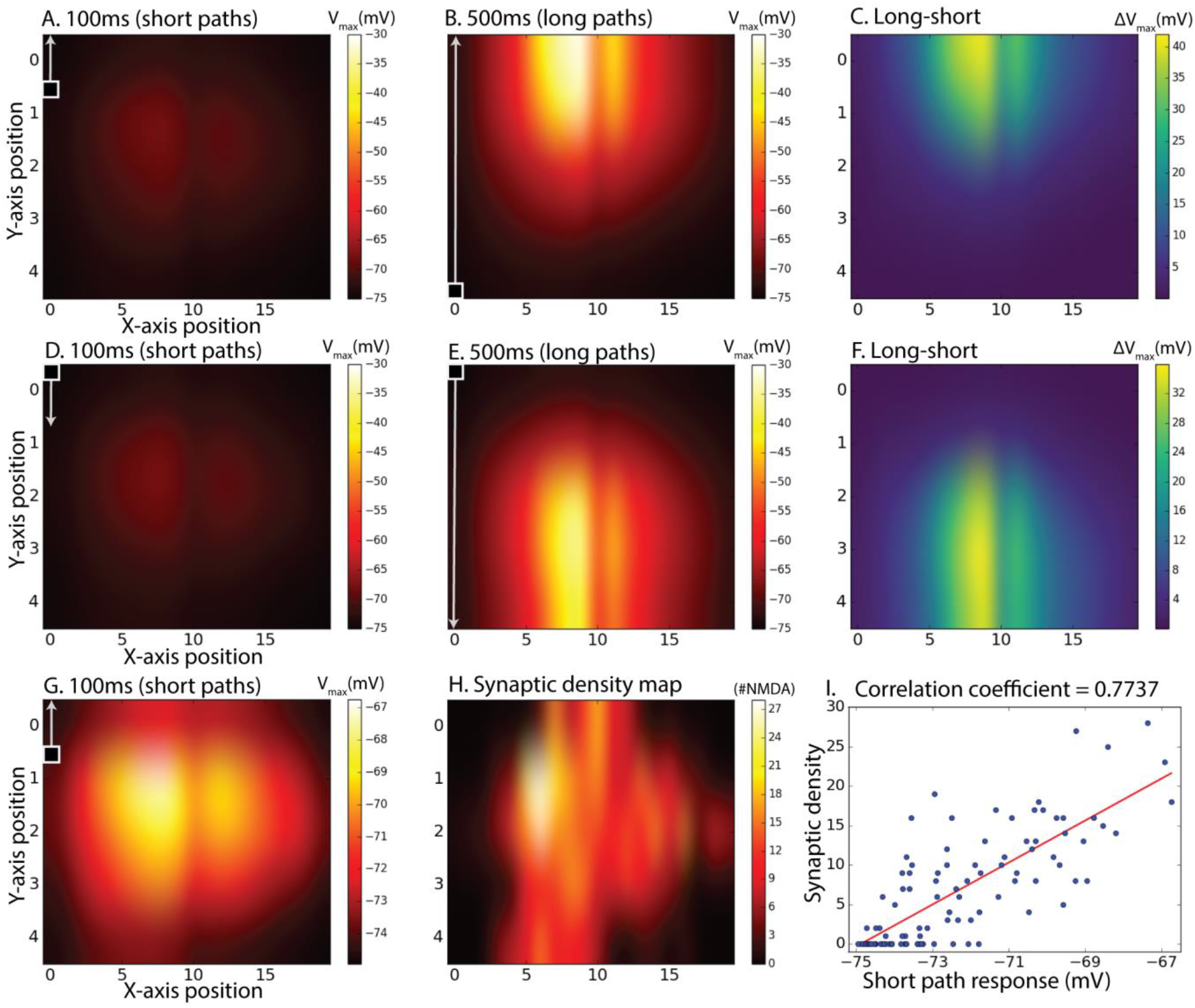
Subdivided long continuous paths versus short paths, difference between them and comparison with synaptic density map. The 100ms measurement period is indicated with a square target with an arrow and also applies to the second row **(D,E,F)**. G_max_=8.25pS was used. **(A)** Maximum membrane potential from short (100ms) paths starting at positions corresponding to a 20 by 5 matrix over the dendritic tree. **(B)** Maximum membrane potential from continuous target paths with responses subdivided to the same paths as in **(A)**. In **(A,B)**, the targets were moving upward. **(C)** Equals difference between **(B)** and **(A)** to achieve a response without response elicited by inhomogeneous synaptic density. **(D,E,F)** is the same as **(A,B,C)** but with downward moving targets instead of upward. **(G)** is the same as **(A)** but with individual color-bar scale. **(H)** is a synaptic density map showing the number of synapses (sections) in the specified area. **(I)** is a scatter plot of **(G,H)**.

## 3 Discussion

We have reported here analysis of NMDA-mediated response facilitation using an elaborated version of a model for which we previously reported the modelling framework (Bekkouche et al., 2017). It should be noted that our preliminary version of this model suffered from mapping inaccuracies between the ESTMD inputs and the BSTMD1 model. Our new model shows facilitation that gradually increases for targets moving on long trajectories (**Figure 7**), as also seen in dragonfly neurons (Nordström et al., 2011; Dunbier et al., 2012) as opposed to a more random or transient looking shapes that we observed previously. Another factor that was removed in this new version of the model is the fact that the ESTMD here only selects one target and the actual output value is not used. This new aspect of the model was introduced to better mimic spiking activity and to remove the clustered response levels that could be seen in (Bekkouche et al., 2017). These clusters emerged due to mismatched mapping between input images and ESTMD array size. Although loss or gain of visual information due to subsampling may be a way for neural networks to generate heterogeneous response from two equally salient stimuli, we decided to not use the actual pixel values in the present study to ignore this effect. To our surprise the clustering persisted, which may still be due to the same effect mediated through the ESTMD to BSTMD1 model activation threshold. When the target is well aligned with an individual ESTMD, the response is stronger and triggers the threshold earlier, compared to a mismatched target to ESTMD case. These differences in triggering time likely have large effects due to NDMAs amplifying properties.

To conclude, our results show that the NMDA synapses enable strongly enhanced responses for continuously moving stimuli, but not at all for random jumps or stimuli moving on short paths.

### 3.1 Dendritic morphology

Our results suggest that the BSTMD1 dendritic morphology may be optimized for responding to continuous target motion. As we have shown, the very dense dendritic field of BSTMD1 includes a larger number of bifurcations, branches and tips than the fly wide-field motion sensitive neuron, HSE. While our analysis strongly suggests that it is this synaptic density that contributes to the large difference we observed in the degree of facilitation in these two models, confirmation of this finding would require more careful investigation of the effect of different dendritic trees on these kinds of synaptic input mapping experiments. Our very detailed BSTDM1 dendritic tree reconstruction certainly provides a basis for such comparative studies and may serve as a useful resource for research groups working on similar hybrid models but who have in the past resorted to using artificial dendritic trees (e.g. Farah et al., 2017) or one from another neuronal subtype (Shoemaker, 2015). This also highlights both the need for, and the challenges in obtaining high-resolution 3-dimensional imagery for complete reconstruction of dendritic trees using state-of-the-art tissue clearing and confocal imaging techniques (Bekkouche et al., 2020).

### 3.2 Model similarity to *in vivo* recordings and limitations

Our model was able to faithfully capture two key aspects of facilitation observed in the real BSTMD1 neuron, i.e. both the time course of the response build-up due to recruitment of nonlinear facilitation and the presence of a localised facilitation ‘spotlight’ which may enhance the intrinsic salience of an attended target. The similarity of facilitation curves to *in vivo* recordings adds to the validity of the model. However, the threshold for the ESTMD model or other model parameters may need to be adjusted in future versions of the model to account for more detailed response pattern differences. A further increment of validity could be achieved by using optimization tools for adapting ion channel parameters based on *in vivo* recordings such as the one shown in **Figure 5A**.

Although this serves as an important confirmation that our particular model framework, using a dendritic network and NMDA synapses, is sufficient to exhibit bio-mimetic facilitation, we cannot expect the primary elaboration of the model - the inclusion of NDMDA synapses - to reproduce the entire repertoire of properties of facilitation as seen in the dragonfly neurons. In particular, facilitation in these neurons has recently been shown to be predictive, such that the locus of facilitation moves forward over time within the visual field, anticipating the future path of the target. Our model lacks any temporal wave propagation process that could explain this behavior. Apart from the spiking mechanism model version, the only currents included in our model have essentially passive properties, and the graded electrical current spreads with such high velocity in the dendrites that any positional information is quickly lost. It is quite likely that calcium channels (apart from NMDA receptors) both in the cell membrane and on the endoplasmic reticulum play a role in regulating facilitation and its spread *in vivo*, and there is clearly scope for future revisions of our model to incorporate that feature, for example.

Another potential way of solving this limitation is to model a network of more local NDMA neuron units that are still able to facilitate locally, but with interactions between the units allowing them to conduct a wave of activity across the network, instead of relying on the dendritic network as an integrator across the whole visual field. This way, the spread of activity relies on the transmission properties of the synapses between the interacting neurons instead of the velocity of current spread across a membrane. Future models using a similar dendritic density to the one that we have shown here could for example incorporate a network of local STMDs, rather than a single large field unit, to simulate the predictive wave of facilitation. The so-called “small-field” STMD neurons described from both dipteran flies (Barnett et al., 2007) and dragonflies (Wiederman et al., 2017) could be suitable candidates for such an interacting retinotopic network. “Small Field” in this context refers to their receptive field size, which is still around 10° in angular subtense: small compared with neurons such as BSTMD1 which respond across the whole visual panorama, but still on the order of 100 or more local inputs (ommatidia) at the level of early visual processing. This is a large enough size to allow both local target selective processing and facilitation. Predictive behavior such as a wave of facilitation could then be implemented either by excitatory and inhibitory interactions between neighboring SF-STMDs, or via an additional wave process implemented within the dendrites of a downstream neuron such as BSTMD1. For example, an untested idea is to incorporate graded NMDA/excitatory receptors which would be able to transfer facilitation without the need of spikes, as seen in vivo in dragonfly CSTMD1 (Wiederman et al., 2017). In addition, the network of neurons could implement the winner-takes-all mechanism and test hypotheses related to selective attention. this would allow testing of, for example, how facilitation is controlled and suppressed by excitation and inhibition to enhance attention towards one target and suppress another, a phenomenon that is challenging to study *in vivo* (Lancer et al., 2019).

### 3.3 Future developments

The results of this study together with ongoing work with single-compartment models will provide the ground work for developing a morphologically and biophysically detailed network model of the primary brain regions involved in small target motion processing. Together with the bioinspired model, this would lead to first model involving the whole optic lobe with such detail. We believe combining highly detailed models with bioinspired models is necessary to push the field of computational neurobiology forward without having to wait for computational capability to increase and all biological details to be discovered or proven, which we may not happen for a long time. We believe the use of these bioinspired models is more reasonable for brain areas which are considered more mechanistic and which we know relatively more about (receptors, lamina, part of the medulla). Using this hybrid model approach, we will be able to build full scale brains a lot sooner than otherwise. A full scale computational brain model would provide a unique environment for testing neural mechanisms and disease states that could potentially lead to improved object tracking systems, new variants of deep learning or machine learning algorithms, or discovery of medically relevant functions of neural networks involved in attention leading to new hypotheses for development of drugs or other treatments.

## 6 Author contributions

Conceptualization: BB, DC, PS. Modelling and simulation: BB. NMDA synapse analysis: BB, PS. Dendritic tree analysis: BB, ER. *In vivo* electrophysiology & filling of BSTMD1: JF, SW. Confocal imaging: DC, JF. BSTMD1 reconstruction: JF, BB. Manuscript writing: BB, DC. Comments on writing: all.

## 7 Acknowledgements

We thank Robert McDougal for providing an example SWC morphology import procedure on the NEURON forum.

## 9 Data Availability Statement

The model is available upon request.

